# The Fe-S cluster assembly factors NFU4 and NFU5 are primarily required for protein lipoylation in mitochondria

**DOI:** 10.1101/2021.09.24.461724

**Authors:** Jonathan Przybyla-Toscano, Andrew E. Maclean, Marina Franceschetti, Daniela Liebsch, Florence Vignols, Olivier Keech, Nicolas Rouhier, Janneke Balk

**Affiliations:** Université de Lorraine, INRAE, IAM, F-54000 Nancy, France; Department of Biological Chemistry, John Innes Centre, Norwich NR4 7UH, UK; School of Biological Sciences, University of East Anglia, Norwich NR4 7TJ, UK; Department of Plant Physiology, Umeå Plant Science Centre, Umeå University, S-90187, Umeå, Sweden; BPMP, Université de Montpellier, CNRS, INRAE, SupAgro, Montpellier, France

**Author notes:** Joint first authors (Jonathan Przybyla-Toscano, Andrew E. Maclean). Wellcome Centre for Integrative Parasitology, University of Glasgow, 120 University Place, Glasgow, G12 8TA, UK. Instituto de Biología Molecular y Celular de Rosario, IBR-CONICET, Ocampo y Esmeralda s/n, 2000 Rosario, Argentina. The authors responsible for distribution of materials integral to the findings presented in this article in accordance with the policy described in the Instruction of Authors are: Janneke Balk and Nicolas Rouhier. Author contributions: N.R. and J.B. conceived the study and the overall research plans; J.P.-T., A.E.M., M.F., D.L., F.V., O.K and J.B. designed and performed the experiments; all authors analysed the data. J.B. wrote the article with figures and text contributions of all authors.

## Abstract

Plants have evolutionarily conserved NFU-domain proteins that are targeted to plastids or mitochondria. The ‘plastid-type’ NFU1, NFU2 and NFU3 in *Arabidopsis thaliana* play a role in iron-sulfur (Fe-S) cluster assembly in this organelle, whereas the type-II NFU4 and NFU5 proteins have not been subjected to mutant studies in any plant species to determine their biological role. Here we confirm that NFU4 and NFU5 are targeted to the mitochondria. The proteins are constitutively produced in all parts of the plant, suggesting a housekeeping function. Double *nfu4 nfu5* knockout mutants were embryonic lethal, and depletion of the proteins led to growth arrest of young seedlings. Biochemical analyses revealed that NFU4 and NFU5 are required for lipoylation of the H proteins of the glycine decarboxylase complex and the E2 subunits of other mitochondrial dehydrogenases, with little impact on Fe-S cluster-containing respiratory complexes and aconitase. Consequently, the Gly-to-Ser ratio was increased in mutant seedlings and early growth was improved by elevated CO_2_. In addition, pyruvate, 2-oxoglutarate and branched-chain amino acids accumulated in the *nfu4 nfu5* mutants, further supporting defects in the other three mitochondrial lipoate-dependent enzyme complexes. NFU4 and NFU5 interacted with mitochondrial lipoyl synthase (LIP1) in yeast 2-hybrid and bimolecular fluorescence complementation assays. These data indicate that NFU4 and NFU5 have a more specific function than previously thought, in providing Fe-S clusters to lipoyl synthase.

**One sentence summary:** A pair of evolutionarily conserved proteins involved in iron-sulfur cofactor assembly have a specific role in lipoate biosynthesis for mitochondrial dehydrogenases.

## INTRODUCTION

Iron-sulfur (Fe-S) clusters are inorganic cofactors with redox or regulatory functions. These labile metal cofactors are essential for the function of more than hundred different proteins involved in electron transfer, catalysis or regulatory processes in plants(Przybyla-Toscano et al., 2021a). The biosynthesis of Fe-S clusters requires dedicated assembly proteins that are conserved from bacteria to eukaryotes. Plants contain three Fe-S cluster assembly pathways, the so-called ISC (iron-sulfur cluster) pathway in the mitochondria, the SUF (sulfur mobilization) pathway in the plastids and the CIA (cytosolic iron-sulfur protein assembly) pathway for the maturation of Fe-S proteins in the cytosol and nucleus (Couturier et al., 2013; Balk and Schaedler, 2014). The mitochondrial ISC pathway provides Fe-S clusters for the respiratory complexes I, II and III of the electron transport chain, aconitase in the tricarboxylic acid cycle and several enzymes in biosynthetic pathways (Przybyla-Toscano et al., 2021).

The proposed working mechanism of the ISC pathway is based on mutant studies in Baker’s yeast, *Saccharomyces cerevisiae*, as well as extensive *in vitro* studies using orthologs from other eukaryotes (Lill and Freibert, 2020; Przybyla-Toscano et al., 2021b). In brief, and using nomenclature of the plant proteins, the first step of Fe-S cluster assembly is carried out by the cysteine desulfurase NFS1, which acquires a protein-bound persulfide (S^0^) from cysteine. The persulfide is transferred to the scaffold protein ISU1, where it is reduced to sulfide (S^2-^) and combined with Fe to form a Fe_2_S_2_ cluster. Next, a chaperone-cochaperone couple helps move the cluster from ISU1 to one of several so-called transfer proteins which facilitate incorporation of the correct type of cluster into client Fe-S proteins. The precise molecular roles of these transfer proteins are not known, and research has focussed on establishing their sequence of action and whether they are required for specific Fe-S proteins. The mitochondrial glutaredoxin GRXS15 is likely to accept the Fe_2_S_2_ cluster from ISU1, as shown for yeast GRX5 (reviewed in Lill and Freibert, 2020). *In vitro* studies demonstrated that Arabidopsis GRXS15 can bind an Fe_2_S_2_ cluster and transfer it directly to ferredoxin 1 (Moseler et al., 2015; Ströher et al., 2016; Azam et al., 2020b). (Moseler et al., 2015; Azam et al., 2020b)(Moseler et al., 2015; Azam et al., 2020b)(Moseler et al., 2015; Azam et al., 2020b)(Moseler et al., 2015; Azam et al., 2020b)(Moseler et al., 2015; Azam et al., 2020b)(Moseler et al., 2015; Azam et al., 2020b)These glutaredoxins also mediate the reductive fusion of two Fe_2_S_2_ clusters to form a Fe_4_S_4_ cluster on the heterodimeric ISCA1/2 proteins (Brancaccio et al., 2014; Azam et al., 2020a; Weiler et al., 2020). Further *in vitro* studies showed that ISCA1/2 can transfer the Fe_4_S_4_ cluster to NFU4 or NFU5, which can donate the cluster to aconitase 2 (Azam et al., 2020a). All mitochondrial NFU proteins studied to date form homodimers with a Fe_4_S_4_ cluster bound to the CxxC motifs of each protomer. The Fe-S cluster binding CxxC motif is also found in INDH (Iron-sulfur cluster protein required for NADH Dehydrogenase) which has been proposed to specifically transfer Fe-S clusters to respiratory complex I (Wydro et al., 2013).

The NFU (NifU-like) proteins are named after the *nifU* gene in the nitrogen fixation (*nif*) regulon in *Azotobacter vinelandii* (Yuvaniyama et al., 2000). In fact, AvNifU has three protein domains that usually exist as separate proteins in other organisms: the N-terminal domain of AvNifU is homologous to ISU1, the central domain is a functional ferredoxin, and the C-terminal domain is similar to what are generically called NFU proteins (Mühlenhoff and Lill, 2000; Py et al., 2012). Based on phylogenetic analysis, four different types of NFU proteins have been identified in bacteria, of which type II in alpha-proteobacteria has been inherited by mitochondria (Py et al., 2012). In the green lineage, NFU proteins are also present in plastids. They fall outside the bacterial classification system and are characterized by a duplicated NFU sequence, of which the second sequence is degenerate and lacks the CxxC motif. Of the five NFU proteins in Arabidopsis, NFU1, NFU2 and NFU3 belong to the ‘plastid-type’ and are indeed targeted to plastids (León et al., 2003). NFU4 (AT3G20970) and NFU5 (AT1G51390) are type-II NFU proteins. Mitochondrial localization of Arabidopsis NFU4 was supported by transient expression of a GFP fusion protein (León et al., 2003). Furthermore, it was shown that *NFU4* and *NFU5* are able to complement the yeast *nfu1*Δ and *nfu1*Δisu*1*Δ strains with respect to growth and biochemical phenotypes (León et al., 2003; Uzarska et al., 2018), demonstrating that the plant genes are functional orthologs of yeast *NFU1*. However, the physiological role of mitochondrial NFU proteins has not been investigated in plants.

Deletion of *NFU1* in yeast impairs growth on acetate as carbon source and affects specific Fe-S enzymes, such as aconitase, respiratory complex II (succinate dehydrogenase, SDH), and also lipoyl-dependent enzymes (Melber et al., 2016). The severity of the *nfu1*Δ phenotype depends on the yeast strain but is relatively mild in contrast to the essential nature of *NFU1* in mammals. A small number of mitochondrial disorders in human patients has been associated with rare mutations in the coding sequence of *NFU1* (Cameron et al., 2011; Navarro-Sastre et al., 2011; Mayr et al., 2014). *NFU1* patients commonly display infantile encephalopathy with symptoms such as hyperglycinaemia and lactic acidosis, which in severe cases is fatal in the first year of life. Biochemical tests on muscle biopsies or fibroblasts showed decreased activities of lipoylated enzymes and, in some but not all cases, a decrease in aconitase and SDH activity.

In eukaryotes, there are four mitochondrial lipoyl-dependent enzyme systems, pyruvate dehydrogenase (PDH), α-ketoglutarate dehydrogenase (KGDH), branched-chain α-ketoacid dehydrogenase (BCKDH) and glycine decarboxylase complex (GDC), also referred to as the glycine cleavage system. The PDH, KGDH and BCKDH complexes consist of three different subunits (E1 - E3), of which E2 is lipoylated. The structure of GDC complex differs, being composed of four proteins named L, P, T and H, with lipoyl bound to the H protein. The lipoyl cofactor (6,8-dithiooctanoic acid) mediates the oxidative decarboxylation reactions carried out by those enzyme complexes (Mayr et al., 2014). The 8-carbon fatty acid is covalently bound to proteins via a lysine residue, forming a lipoamide, and contains two sulfur atoms at C6 and C8 that are inserted by lipoyl synthase. The sulfur atoms are provided by the auxiliary Fe_4_S_4_ cluster of lipoyl synthase. Therefore, this cluster needs to be reassembled after every catalytic cycle. *In vitro* studies showed that the bacterial NfuA protein enabled catalytic rates of lipoyl formation by its ability to reconstitute the auxiliary cluster of lipoyl synthase (McCarthy and Booker, 2017).

Here we show that the Arabidopsis NFU4 and NFU5 proteins perform an essential function in mitochondria. Analysis of several mutant lines showed that low levels of the NFU proteins are sufficient for normal growth, but their near-complete depletion revealed a defect in protein lipoylation, affecting substrate turnover by lipoate-dependent enzyme complexes such as GDC, KGDH and BCKDH.

## RESULTS

### NFU4 and NFU5 are soluble mitochondrial matrix proteins

A previous study showed that transient expression of a fusion protein of the first 116 amino acids of NFU4 with GFP was targeted to mitochondria in Arabidopsis protoplasts (León et al., 2003). However, the localization of NFU5 was not experimentally tested. Some evidence for the presence of NFU5 in mitochondria has been obtained by proteomics studies (Ito et al., 2006; Tan et al., 2010; Fuchs et al., 2020), but has so far not been confirmed by other approaches. Therefore, we cloned the predicted mitochondrial targeting sequences (MTS) of *NFU4* and *NFU5* in frame with the *GFP* coding sequence. The length of the MTS of each protein was determined by combining several *in silico* analyses such as target peptide prediction algorithms, the distribution of acidic amino acids and alignment with eukaryotic and prokaryotic homologs. Following this, amino acids 1 to 79 for NFU4 and 1 to 74 for NFU5 were assigned as MTS. The fusion genes were placed under the control of a double *CaMV 35S* promoter and transformed into Arabidopsis protoplasts. Both NFU4_MTS_-GFP and NFU5_MTS_-GFP showed a punctate pattern of GFP which co-localized with MitoTracker dye, but was distinct from the chloroplast autofluorescence (Supplemental Fig. S1A). Protein blot analysis of total and mitochondrial protein fractions confirmed the enrichment of the native NFU4 and NFU5 proteins in mitochondria (Supplemental Fig. S1B). Furthermore, fractionation of the mitochondria into soluble matrix and membrane fractions indicated that NFU4 and NFU5 are matrix proteins, similar to ISU1 and GRXS15 (Supplemental Fig. S1C).

### NFU4 and NFU5 proteins are abundant in all plant organs

Antibodies raised against NFU4 cross-reacted with NFU5 and vice versa, which is not surprising as the mature proteins share 90% amino acid identity. To determine which of the two immune signals with similar gel mobilities belongs to NFU4 and NFU5, and for subsequent mutant studies, we obtained three mutant alleles each for *nfu4* and *nfu5* (Fig. 1A).

**Figure 1.**
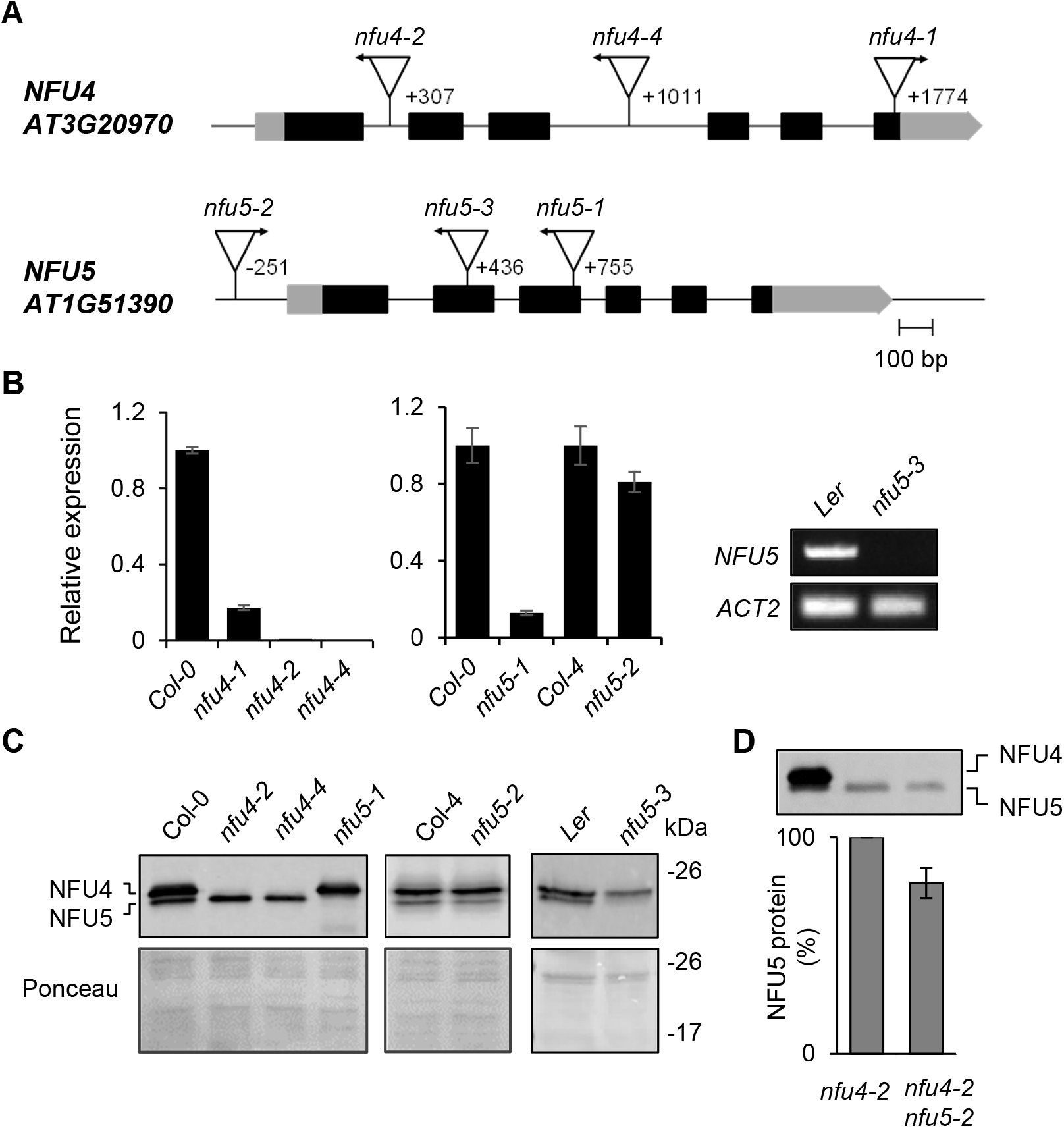
Genetic analysis of Arabidopsis mutants in *NFU4* and *NFU5*. **A.** Gene models of *NFU4* and *NFU5* and the positions of T-DNA insertions. Black bars represent exons, grey bars are the 5’ and 3’ untranslated regions of the transcript. Triangles represent T-DNA insertions, their orientation is marked with an arrow to indicate the outward facing left border primer. The position of the T-DNA relative to the ATG start codon is indicated by the number of the nucleotide next to the left-border sequence. **B.** Transcript levels of *NFU4* and *NFU5* in leaf tissue of wild type (Col-0, Col-4 or Ler) and the indicated T-DNA insertion lines, determined by RT-qPCR (graphs) or standard RT-PCR (right). For RT-qPCR, values are the average of 3 biological samples ± SE. **C.** Protein blot analysis of NFU4 and NFU5 in mitochondria isolated from seedlings. Blots were labelled with antibodies against NFU4. Ponceau S stain was used to confirm equal loading and transfer. **D.** Decrease in NFU5 protein as a consequence of the *nfu5-2* allele, quantified in the *nfu4-2* mutant background. See Supplemental Fig. S2 for more details of the quantification.

Quantitative reverse transcription PCR (RT-qPCR) showed a virtual absence of *NFU4* transcripts in homozygous *nfu4-2* and *nfu4-4* single mutant plants, whereas ~10% transcript remained in the *nfu4-1* mutant (Fig. 1B). Expression of *NFU5* was strongly diminished in the *nfu5-1* line (Fig. 1B). The *nfu5-2* mutant had approximately 15% less *NFU5* transcript compared to its respective wild type, consistent with insertion of the T-DNA in the promoter, 251 nucleotides upstream of the ATG start codon. For the *nfu5-3* mutant, qPCR analysis suggested that the transcript levels of *NFU5* were strongly increased, but this result is likely due to the presence of an outward facing promoter sequence at the right T-DNA border and the qPCR primers being downstream of this. Probing for full-length *NFU5* transcript by RT-PCR showed that the transcript is lacking in the *nfu5-3* mutant (Fig. 1B, right panel), as expected from the position of the T-DNA in exon 2 (Fig. 1A).

Protein blot analysis using anti-NFU4 serum showed that the upper immune signal is absent in the *nfu4-2* and *nfu4-4* mutants, and that the lower signal is missing in the *nfu5-1* and *nfu5-3* mutant alleles (Fig. 1C). The native protein products of NFU4 and NFU5 could thus be assigned and their electrophoretic mobilities match the calculated molecular weights of 22.1 kDa for NFU4 and 21.7 kDa for NFU5 without their predicted MTS. The levels of NFU5 protein produced from the *nfu5-2* allele were assessed by semi-quantitative protein blot analysis in plants homozygous for the *nfu4-2* allele, to rule out signal contribution from NFU4. The NFU5 protein level was ~15% less in the *nfu4-2 nfu5-2* double mutant compared to wild type (Fig.1D; Supplemental Fig. S2), matching the decrease in *NFU5* transcripts (Fig. 1B).

Publicly available RNA-seq data (bar.utoronto.ca) indicated that *NFU4* and *NFU5* are expressed throughout the plant’s life cycle and in all organs. Protein blot analysis using protein extracts from roots, leaves, stems and reproductive organs confirmed that NFU4 and NFU5 proteins are expressed in all organs, with slightly lower levels in stems and slightly higher levels in flower buds and flowers, based on total protein (Fig. 2A). Densitometry and quantification of the immune signals relative to known amounts of recombinant protein indicated that NFU4 was approximately 2-fold more abundant than NFU5 (Fig. 2B), in agreement with a recent quantitative analysis of all mitochondrial proteins (Fuchs et al., 2020).

**Figure 2.**
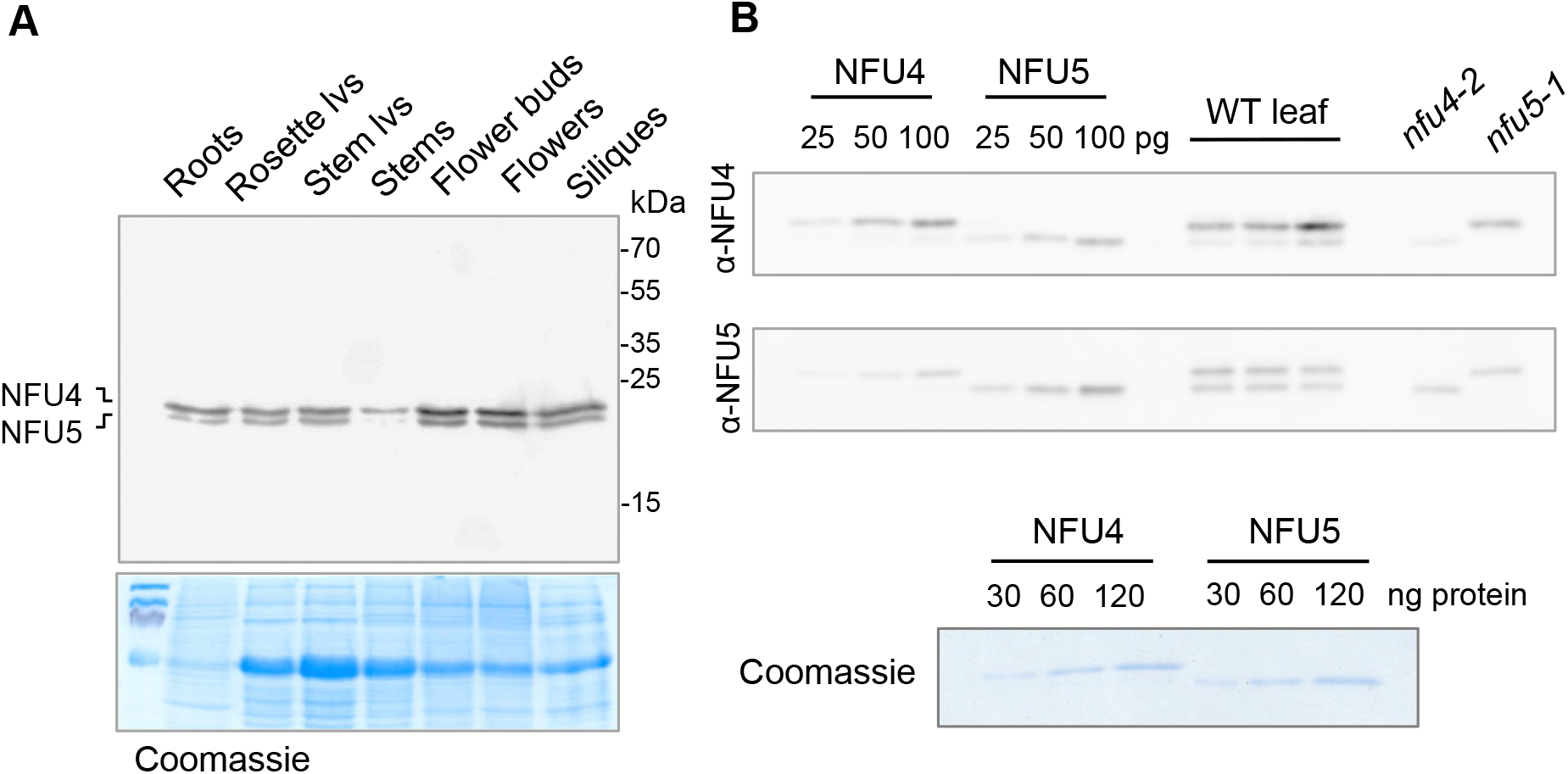
NFU4 and NFU5 proteins are abundant in all plant organs. **A.** Protein blot analysis of NFU4 and NFU5 in different organs of a 6-week-old Arabidopsis plant (Col-0), 20 μg protein per lane, labelled with NFU4 antibodies. Coomassie Blue staining of the gel after transfer was used as loading control. lvs, leaves. **B.** Specific affinity of the polyclonal antibodies raised against NFU4 and NFU5. Luminescence signals of known amounts of recombinant proteins were compared with signals in purified mitochondria from wild-type (WT) leaves and from cell culture of *nfu4-2* and *nfu5-1* mutants. Each antiserum cross-reacts with the other isoform (90% amino acid identity), but has a stronger affinity for the protein it was raised against.

### *NFU4* and *NFU5* are redundant genes, but together are essential for embryo development

In order to evaluate the physiological importance of NFU4 and NFU5, the available T-DNA insertion lines were grown on soil under long-day conditions. The three different mutant lines for *NFU4* or *NFU5* showed normal growth and development of the rosette leaves compared to their respective wild-type plants (Fig. 3A). The growth rate of primary roots in young seedlings, which is particularly sensitive to mitochondrial defects, was decreased by 8% in the *nfu4-2* mutant but a decrease in the *nfu4-4* mutant was not statistically significant (Supplemental Fig. S3A, B). Root growth in the *nfu5-1* mutant was ~30% decreased, whereas *nfu5-2* root length was similar to wild type (Supplemental Fig. S3C, D), as expected based on the weak nature of this mutant allele. In addition, the seedlings were challenged with Paraquat (methyl viologen) in the medium, which strongly inhibited growth of an *E. coli nfuA*Δ strain (Angelini et al., 2008) and is a well-known inducer of oxidative stress in mitochondria (Cochemé and Murphy, 2008). A concentration of 10 nM Paraquat was chosen based on experimental calibration, resulting in 20% inhibition of wild-type root growth. However, Paraquat treatment did not specifically affect the mutants more than the wild-type controls (Supplemental Fig. S3). Taken together, these observations indicate that aside from a role in root elongation, individually neither NFU4 nor NFU5 perform a critical function under normal growth conditions.

**Figure 3.**
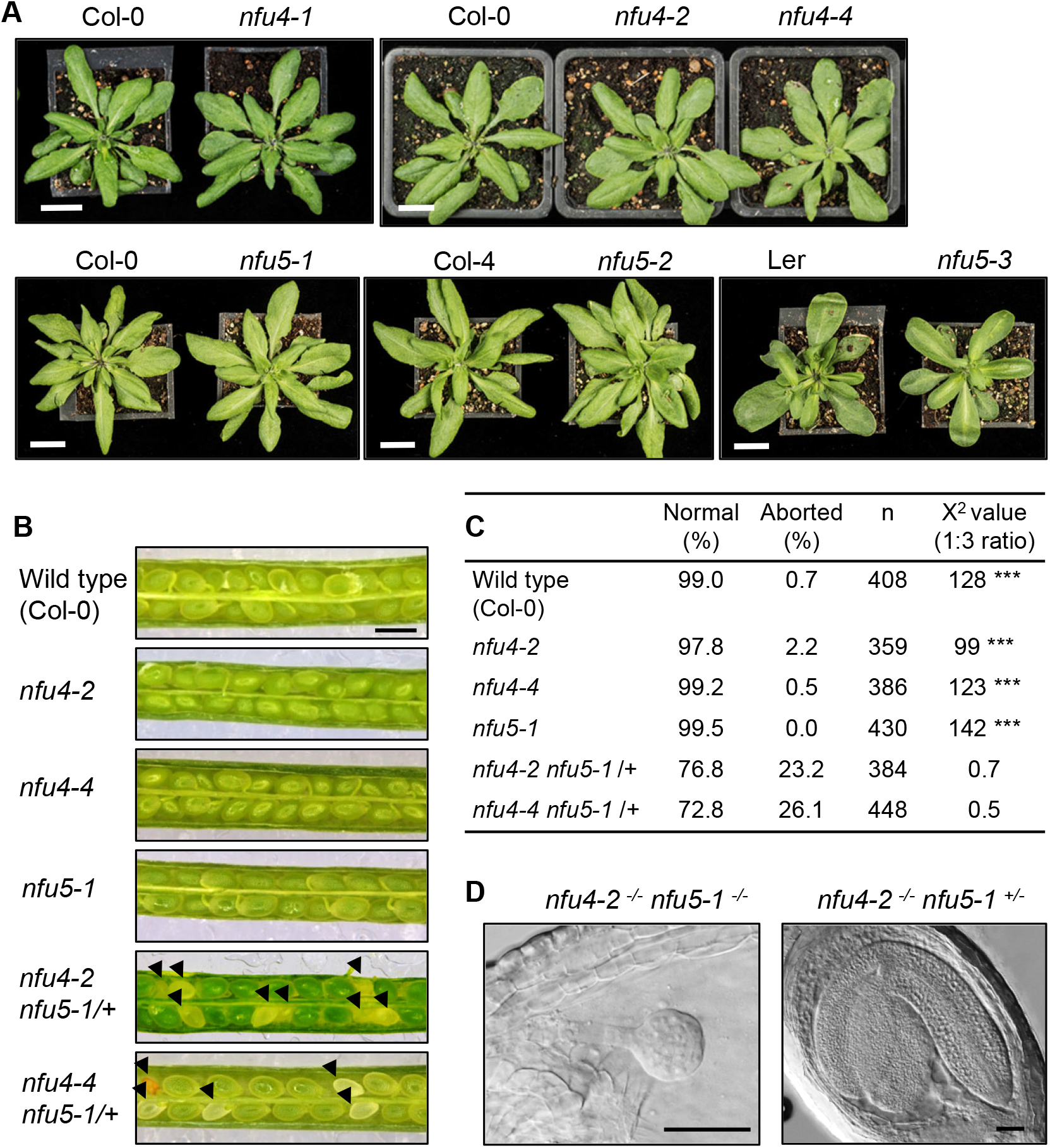
Phenotypes of *nfu4* and *nfu5* single and double mutants. **A.** Growth phenotype of 4-week-old plants of the indicated genotype. Scale bar: 1 cm. **B.** Images of open siliques with immature seeds in wild type (Col-0) and the indicated mutant lines. Scale bars: 0.5 mm. **C.** Frequency of normal and aborted embryos in *nfu4 nfu5-1/+* plants. \*\*\**p*<0.0001 for 1:3 segregation ratio (χ^2^ test). **D.** An aborted and healthy embryo from the silique of a *nfu4-2 nfu5-1/+* plant. Plant tissue was cleared with Hoyer’s solution and imaged with DIC microscopy. Scale bars: 50 μm.

To generate double mutants, we made reciprocal crosses between *nfu5-1* and *nfu4-2* or *nfu4-4* plants. In the F2 generation, double mutants could not be isolated despite extensive screening by PCR. Therefore, the siliques of *nfu4-2^-/-^nfu5-1*^-/+^ and *nfu4-4^-/-^nfu5-1*^-/+^ plants were dissected to analyse embryo development. Approximately one quarter of the immature seeds were white (Fig. 3B, C), containing embryos arrested at the globular stage (Fig. 3D). Based on these numbers it is reasonable to assume that the white seeds are double knockout *nfu4 nfu5*, and that a complete lack of both NFU4 and NFU5 protein is detrimental for embryo development.

### Functional relationship between *ISU1*, *NFU4* and *NFU5*

In yeast, the mild growth defect as a result of *NFU1* deletion (*nfu1*Δ) is enhanced in an *isu1*Δ*nfu1*Δ double mutant, underscoring the functional relationship of the ISU1 and NFU1 proteins (Schilke et al., 1999). It should be noted that the yeast *isu1*Δ mutant is viable because of a second *ISU* gene, *ISU2*, but evidently this cannot fully replace the function of *ISU1*. Arabidopsis has three *ISU* paralogs, each of which can functionally substitute for the yeast *ISU1* homolog in the *isu1*Δ*nfu1*Δ strain (León et al., 2005). The *Arabidopsis ISU1* gene is thought to be the main Fe-S scaffold protein in mitochondria, because *ISU2* and *ISU3* have very low expression levels and are almost exclusively expressed in pollen grains (León et al., 2005; Frazzon et al., 2007). In agreement with this, we found that the *isu1-2* allele, in which 61 nucleotides directly upstream of the ATG are deleted due to a T-DNA insertion, is embryonic lethal (Supplemental Table S1). The previously reported *isu1-1* allele, with a T-DNA insertion at nucleotide −65 but with intact sequence up to the start codon, is viable (Frazzon et al., 2007) (Supplemental Fig. S4A). ISU1 protein levels in the *isu1-1* line are decreased to ~20% of wild type (Supplemental Fig. S4B). In cell culture generated from the roots of *isu1-1* seedlings, the levels of NFU4, NFU5, INDH and aconitase were normal, and the activity of respiratory complex I was also comparable to wild type (Supplemental Fig. S4B, C). Interestingly, GRXS15 protein levels were strongly decreased in the *isu1-1* mutant.

Rosettes of *isu1-1* plants were phenotypically indistinguishable from wild type, but seedlings had a 20% decrease in root elongation, which was exacerbated to 40% impairment in the presence of Paraquat (Supplemental Fig. S4D, E). This phenotype may be correlated with decreased GRXS15 levels, since *grxs15* mutants have strongly impaired root growth (Ströher et al., 2016). To investigate genetic interactions between *ISU1* and *NFU4* or *NFU5*, the *isu1-1* line was crossed with *nfu4* and *nfu5* mutant alleles. Crosses involving any of the *nfu5* alleles and *isu1-1* were unsuccessful, despite multiple attempts in reciprocal combinations. In contrast, double *isu1-1 nfu4* mutants were isolated from the F2, and had a shorter root length but not an enhanced growth defect (Supplemental Fig. S4D).

Thus, mild growth impairment of the primary root in seedlings with less than 20% ISU1 is not enhanced by deletion of *NFU4*. This is in agreement with the model that NFU proteins function downstream from ISU1 in Fe-S cluster transfer, and not as an alternative assembly scaffold.

### Low amounts of NFU5 alone are sufficient for normal mitochondrial function in vegetative tissues

The normal rosette growth of single *nfu4* and *nfu5* mutants does not preclude defects in Fe-S enzyme activities. For example, up to 80% of complex I activity can be lost in Arabidopsis without major growth penalties (Meyer et al., 2011). To analyse the levels and/or activities of major ISC maturation factors and mitochondrial Fe-S enzymes as a consequence of loss of NFU4 and/or NFU5, mitochondria were purified from 2-week-old seedlings. Protein blot analyses showed no differences in the levels of ISU1, GRXS15, INDH and aconitase between *nfu4-2*, *nfu4-4* and *nfu5-1* and wild type (Supplemental Fig. S5A). Labelling with antibodies against lipoate also showed no differences in the abundance of lipoylated proteins (Supplemental Fig. S5B). To analyse the levels of mitochondrial respiratory complexes, we generated cell cultures from primary roots, then purified mitochondria for blue-native PAGE. Protein complexes were stained with Coomassie Blue (Supplemental Fig. S5C), and complex I and complex II were additionally visualized by in-gel activity staining (Supplemental Fig. S5D, E). Again, no differences in the levels or the activities of these major Fe-S cluster-dependent complexes were detected in single mutants compared with wild-type plants.

To obtain viable plants with strongly diminished NFU4 and NFU5 protein levels, mutant alleles were combined via crossing. *nfu4-2 nfu5-2* plants lack NFU4 protein and have ~85% NFU5 compared to wild type, whereas hemizygous *nfu4-2 nfu5-1/nfu5-2* plants have only ~42.5% NFU5 (Fig. 4A). The size and fresh weight of the rosettes of hemizygous plants were similar to wild type (Fig. 4B; Supplemental Fig. S6). In root growth assays, the double mutant seedlings performed similarly to the Col-4 parent. However, in the presence of Paraquat, there was a small but significant decrease in root growth in the hemizygous mutant (Supplemental Fig. S3E). Because of a lipoate biosynthesis defect in yeast and mammalian *nfu* mutants, we further tested growth of the Arabidopsis *nfu4-2 nfu5-2* double mutants under low CO_2_ (150 ppm) which should bring out defects in GDC. However, no obvious differences in either fresh or dry weight biomass were observed between mutant and wild-type plants (Supplemental Fig. S6).

**Figure 4.**
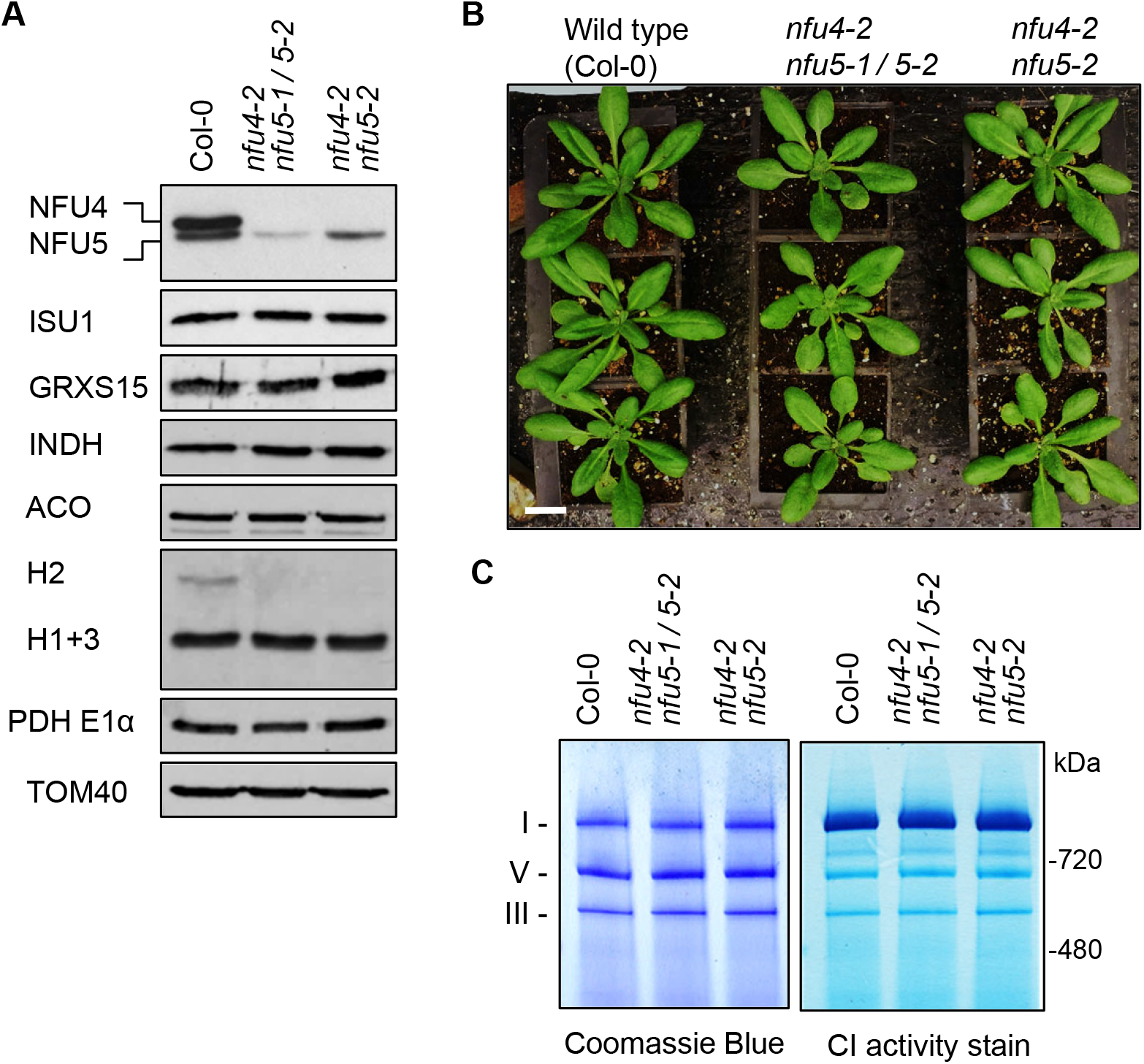
Analysis of *nfu4 nfu5* hemizygous and double mutants. **A.** Protein blot analysis using protein extracts of mitochondria isolated from callus of wild type (Col-0), hemizygous and *nfu4-2 nfu5-2* double mutants as indicated. Antibodies against the following proteins were used: NFU4 and NFU5; the Fe-S scaffold protein ISU1, glutaredoxin GRXS15, complex I assembly factor INDH, aconitase (ACO), the H protein subunit of the glycine decarboxylase complex (GDC), E1α subunit of pyruvate dehydrogenase (PDH) and the translocase of the outer membrane TOM40. **B.** Growth phenotype of 4-week-old wild-type and *nfu4-2 nfu5* plants. Scale bar: 0.5 cm **C.** Blue-Native PAGE of mitochondrial complexes I, III and V stained with Coomassie Blue (left panel) and by NADH/NBT activity staining for complex I (right panel) in the indicated plant lines.

To follow up the growth assays, biochemical analyses were carried out on mitochondria purified from cell culture of hemizygous *nfu4-2 nfu5-1*/*nfu5-2* and homozygous *nfu4-2 nfu5-2* seedlings. Protein blot analyses showed no differences in the levels of the mitochondrial Fe-S cluster assembly proteins ISU1, GRXS15 and INDH and the Fe-S enzyme aconitase (Fig. 4A). The only observed difference was a decrease in the levels of the H2 protein of GDC. The levels of respiratory complexes I and III in the viable *nfu4 nfu5* double mutants were comparable to wild type (Fig. 4C). Taken together, these biochemical studies show that NFU4 and NFU5 are largely redundant during vegetative development; and suggest that small amounts of NFU5 are sufficient to sustain key mitochondrial functions.

### Depletion of NFU4 and NFU5 causes a pleiotropic growth defect and accumulation of substrates of lipoate-dependent enzyme complexes

To uncover the phenotypic effects of a complete lack of NFU4 and NFU5 in vegetative tissues and circumvent embryo lethality, *NFU4* was placed under the control of the *ABI3* promoter (Rohde et al., 1999; Despres et al., 2001) and transformed into hemizygous *nfu4-2*^-/-^ *nfu5-1*/*nfu5-2* plants (Fig. 5A). The *ABI3* promoter drives expression in developing and germinating seeds and can be used to study essential genes (Despres et al., 2001). Positive transformants were selected on hygromycin-containing plates, and three seedlings carrying the *nfu5-1* knockout allele were identified by PCR for further study (independent lines 1, 4 and 10). In the next generation (T2), seedlings segregated in approximately 21 – 25% with a severe growth phenotype and 75 – 79% with normal growth. Leaves of the mutant seedlings turned white upon emergence, and growth was arrested at the second pair of true leaves (Fig. 5B). PCR analysis confirmed that the mutant (m) seedlings lacked a functional copy of *NFU5*, whereas the wild-type-like (wtl) siblings carried the *nfu5-2* allele with an intact *NFU5* open reading frame (Fig. 5C). In agreement with the genotyping results, the mutant seedlings lacked NFU5 protein, whereas the NFU5-specific protein band was present in the wtl seedlings (Fig. 5D). In both mutant and wtl segregants, the protein levels of NFU4 were either undetectable or very low, reflecting the combined effect of the *nfu4-2* allele and *ABI3* promoter-driven *NFU4* expression.

**Figure 5.**
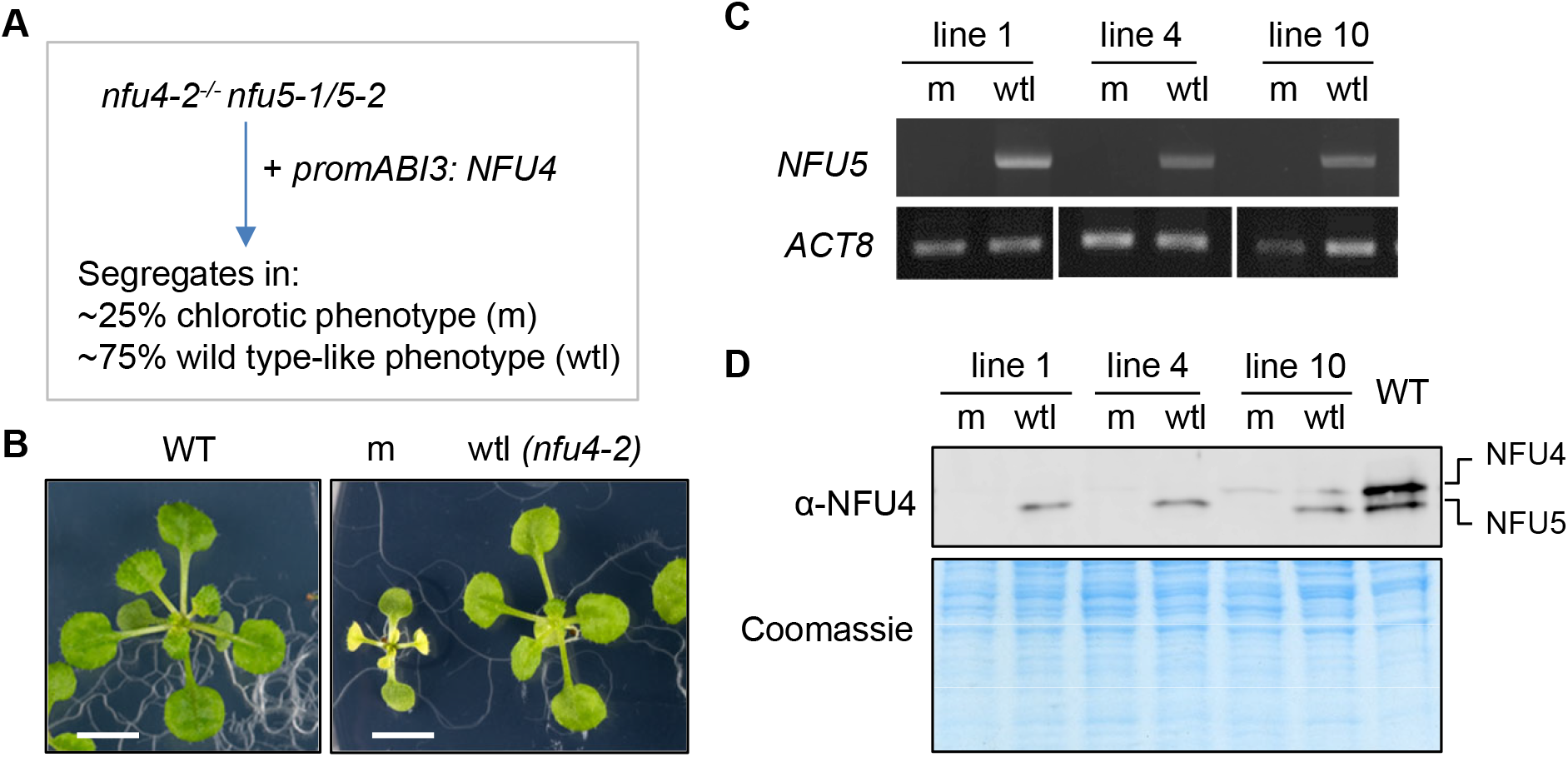
Seedlings depleted of NFU4 and NFU5 proteins have a pleiotropic phenotype. **A.** Scheme for obtaining a mutant line expressing *NFU4* under the control of the *ABI3* promoter in a *nfu4 nfu5* knockout background. The *ABI3* promoter is active during embryogenesis but switched off after seed germination. The observed segregation numbers in T2 seedling from 3 independent lines were: 76 chlorotic/small and 260 wild-type appearance (total n = 336). **B.** Representative images of a wild-type (WT) seedling and the two classes of segregants, mutant (m) and wild type-like (wtl), grown for 21 days on ½ MS medium in 8 h light / 16 h dark cycles. Scale bars: 0.5 cm. **C.** PCR genotyping results of mutant (m) and wild-type-like (wtl) seedlings from 3 independent lines, showing the absence or presence of a functional NFU5 sequence using primers AM84 and AM85. **D.** Protein blot analysis of NFU4 and NFU5 in plants lines as in (C).

In an attempt to overcome growth arrest, the segregating T2 generation was grown under elevated CO_2_. We reasoned that the observed photobleaching of the mutant seedlings could be due to a defect in photorespiration. Under high CO_2_, photorespiration is prevented and impairment of GDC is tolerated, except when it is so low as to affect one-carbon metabolism (Peterhansel et al., 2010). Indeed, when seedlings were germinated and grown under high CO_2_, segregating *nfu4 nfu5* mutant seedlings were more difficult to distinguish from wtl siblings in the first two weeks, as double mutants remained green and started to develop a third pair of true leaves (Fig. 6A). However, soon after transferring the seedlings from agar plates to soil in the high CO_2_ cabinet, the *nfu4 nfu5* mutants turned white and their growth was arrested before the inflorescence was formed (Supplemental Fig. S7).

**Figure 6.**
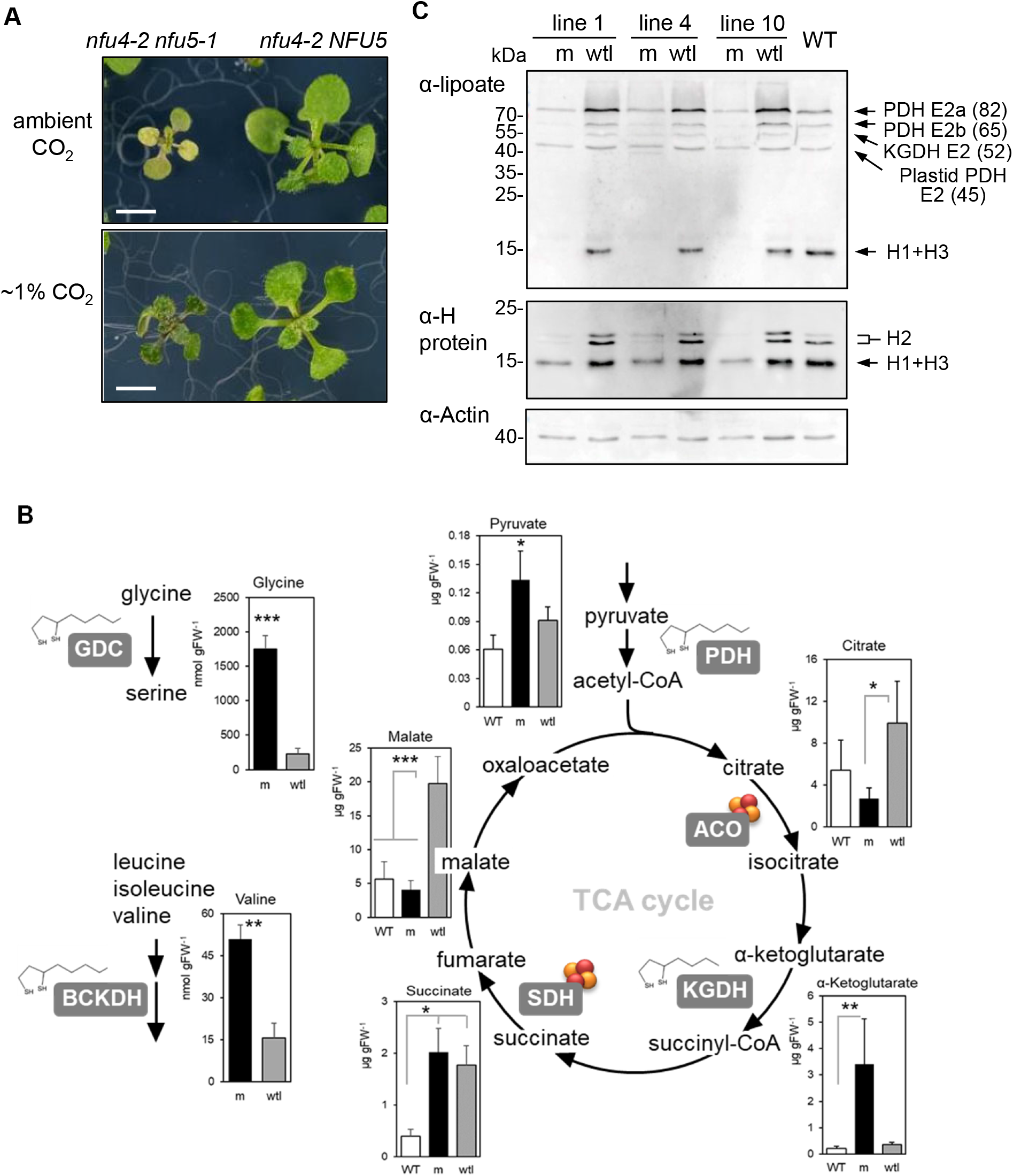
The *nfu4 nfu5* double mutant shows decreased protein lipoylation affecting lipoyl-dependent metabolism. **A.** Representative images of *nfu4 nfu5* double mutant segregants (m) and sibling wild-type like (wtl) seedlings, grown on 1/2 MS agar plates under ambient CO_2_ and 1% CO_2_ in the greenhouse (8/16 h light/dark cycles, variable temperature, ~90% humidity). Scale bars: 0.5 cm. Additional images in Supplemental Fig. S7. **B.** Concentrations of selected free amino acids and organic acids in 3-week-old seedlings of the indicated genotypes. Values represent the average of 3-6 biological replicates ± SD. *p < 0.05, **p < 0.01, ***p < 0.001 (Student *t*-test, pairwise comparison to wild type). See Supplemental Tables S2 and S3 for complete data sets. **C.** Protein blot analysis for lipoyl cofactor (top) and H protein isoforms of glycine decarboxylase complex (bottom), in wild type (WT), *nfu4 nfu5* (m) and wtl segregants. See also Supplemental Fig. S8.

To investigate if GDC activity was decreased in the mutants, we measured the concentration of glycine and serine as well as other free amino acids using LC-MS. Glycine accumulated 8-fold in the mutant compared to wtl seedlings (Fig. 6B; Supplemental Table S2). The Gly:Ser ratio of 3.5 in the mutant versus a ratio of 1 in wtl is indicative of impaired GDC activity in the mutant (Timm et al., 2012; Reinholdt et al., 2021). It should be noted that the serine levels we measured in control (wtl) seedlings were approximately 2-fold lower than routinely measured in rosette leaves of Arabidopsis but the levels of other amino acids were comparable to published values (reviewed in (Hildebrandt et al., 2015)).

Additionally, the branched-chain amino acids leucine and valine were significantly increased in concentration compared with wtl, and the isoleucine concentration was elevated although not significant (Supplemental Table S2). This suggests that the lipoyl-dependent enzyme complex BCKDH has decreased functionality. Of the other amino acids, alanine and phenylalanine levels were on average more than 7-fold increased in the mutant, but the levels were very variable in the 3 biological replicates of the mutant. Similarly, arginine, asparagine, lysine and tryptophan were >2-fold increased in the mutant, and aspartic acid was >2-fold decreased, but not statistically significant because of large variation.

To identify changes in pyruvate and TCA cycle intermediates, we measured the concentrations of selected organic acids by LC-MS/MS. The *nfu4 nfu5* mutant accumulated 10-fold more α-ketoglutarate than the wtl segregants and wild type, indicating that the activity of KGDH is impaired (Fig. 6B; Supplemental Table S3). The concentration of pyruvate in the mutant was 2-fold higher than in wild type seedlings and 1.5-fold higher than in wtl. Citrate and malate concentrations were lower in the *nfu4 nfu5* mutant seedlings, whereas succinate was elevated in both the mutant and wtl segregant compared to wild-type seedlings (Fig. 6B; Supplemental Table S3). Together, these results indicate that NFU4 and NFU5 are required for the function of GDC as well as the other major mitochondrial enzyme complexes that depend on lipoate as a cofactor.

### NFU4 and NFU5 are required for lipoylation of GDC H proteins and E2 subunits of mitochondrial dehydrogenase enzyme complexes

To investigate a possible decrease in protein-bound lipoyl cofactor in the *nfu4 nfu5* mutant, immunoblots of total plant protein extracts were labelled with antibodies against lipoate. This showed a pattern similar to those previously published and proteins were assigned accordingly (Ewald et al., 2014; Ströher et al., 2016; Guan et al., 2017). We observed that *nfu4 nfu5* mutant seedlings lacked a signal with an electrophoretic mobility of 15 kDa corresponding to the lipoylated H1 + H3 proteins of GDC (Fig. 6C, upper panel). The mutants also showed a decreased signal at ~65 kDa, assigned as the E2b subunit isoform of the PDH complex, compared to wild type (WT) and wild-type like (wtl) segregants. The signal corresponding to the 82-kDa E2a subunit of PDH was slightly decreased in the mutant compared to WT, but increased in wtl seedlings. The 45 kDa band was assigned to the E2 subunit proteins of plastid-localized PDH, which had similar levels of lipoylation in the mutant and WT plants.

To probe whether non-lipoylated H proteins were present in the mutant lines, a parallel blot of the same samples was probed with an antibody that recognizes H isoforms from a range of plant species including the 3 isoforms in Arabidopsis (Ströher et al., 2016). The signal from the H1 and H3 isoforms at 15 kDa was strongly decreased in the *nfu4 nfu5* mutant, pointing to destabilization of these subunits due to a lack of bound lipoate. The mature H2 protein has a theoretical mass similar to H1 and H3, but as noted before, it has a lower electrophoretic mobility in SDS-PAGE (Fig. 6C, middle panel), possibly because of its unusual low pI (Lee et al., 2008; Ströher et al., 2016). The two signals assigned as H2 likely correspond to the lipoylated and non-lipoylated forms (Ströher et al., 2016). The major H2 isoform present in wtl and WT was barely detectable in *nfu4 nfu5* seedlings. These data indicate that NFU4 and NFU5 are required for protein lipoylation in mitochondria, strongly affecting the H subunits of GDC and to a lesser extent the E2 subunits of PDH.

### Aconitase and complex II activities are normal in seedlings depleted of NFU4 and NFU5

Since high CO_2_ conditions only partially rescued the severe phenotype of *nfu4 nfu5* mutants, we assessed the overall mitochondrial respiration rate by measuring O_2_ consumption in intact mutant and wtl seedlings in the dark. The *nfu4 nfu5* double mutant showed a 5-fold decrease in respiration compared to wild-type seedlings and segregating wtl siblings (Fig. 7A), in line with defects in key mitochondrial enzyme complexes. In the yeast *nfu1*Δ mutant, the activities of the Fe-S enzymes aconitase and SDH/complex II were decreased by ~25% (Melber et al., 2016; Uzarska et al, 2018). Measurements of aconitase activity in total plant extracts showed a decrease of ~50% in the *nfu4 nfu5* mutant compared to wild-type seedlings and a decrease of 30% compared to wtl segregants, although the latter was not statistically significant (Fig. 7B). However, in plants grown under high CO_2_, aconitase activity was 2-fold higher than under ambient CO_2_ levels and similar in the mutant and wtl segregants (Fig. 7B). This suggests that any decrease in aconitase activity may be a secondary defect, caused by, for example, limited carbon flux through the tricarboxylic acid cycle. This is in agreement with the observed decrease in citrate and malate (Fig. 6B).

**Figure 7.**
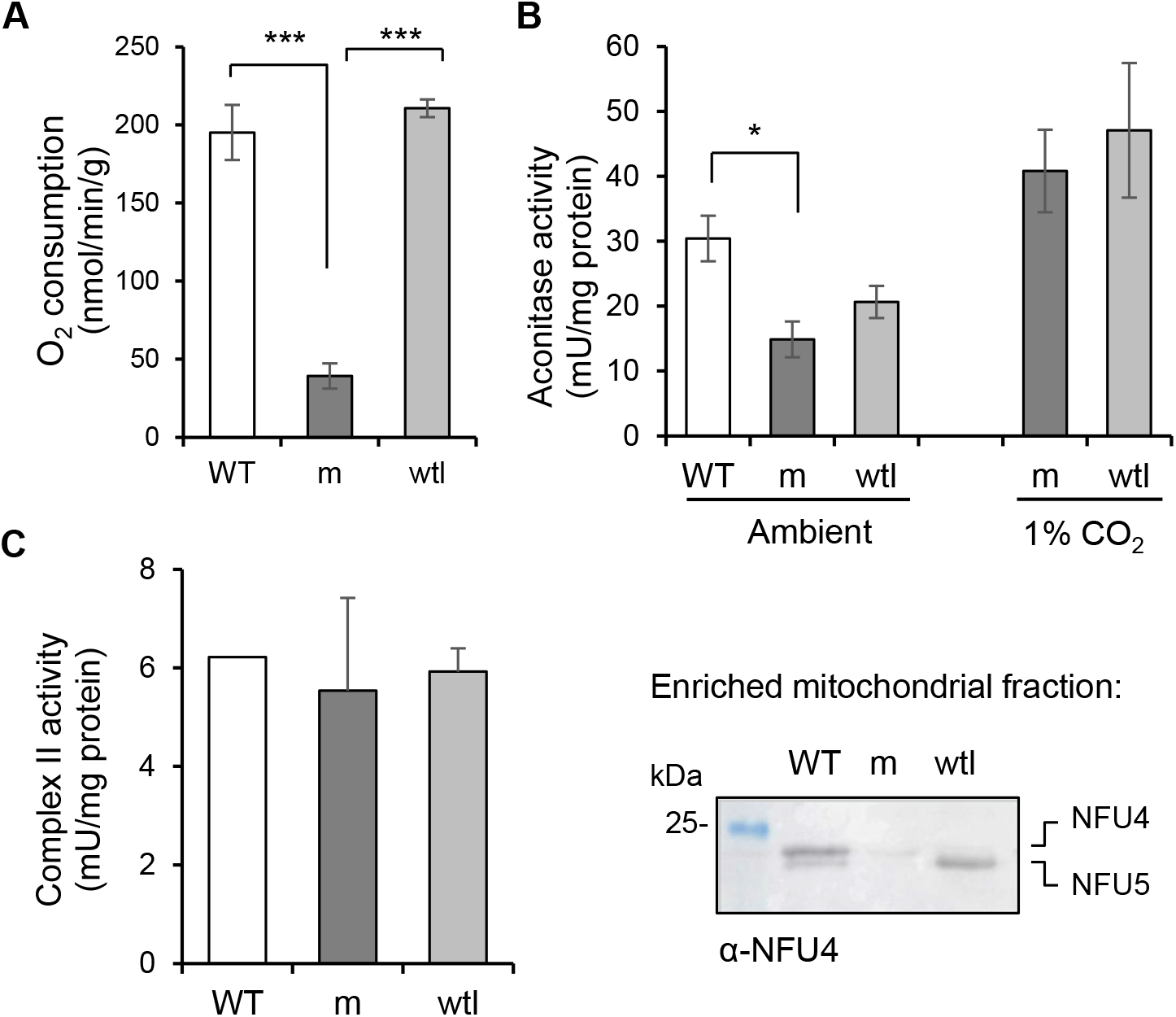
NFU4 and NFU5 are not required for the Fe-S enzymes aconitase and complex II. **A.** Respiration in intact seedlings measured using a liquid-phase oxygen electrode. WT, wild type (Col-0); m, *nfu4-2 nfu5-1* mutant expressing *ABI3prom:NFU4*; wtl, wild-type-like *nfu4-2 NFU5* segregants. Values represent the mean oxygen consumption per g fresh weight ± SD (n = 3). \*\*\**p* < 0.001 (Student *t*-test). **B.** Aconitase activity in total cell extracts in seedlings grown under ambient and 1% CO_2_ for 3 weeks. Values represent the mean ± SD (n = 3-4). \**p* < 0.05 (Student *t*-test). **C.** Complex II activity measured as electron transfer from succinate to ubiquinol (SQR) using the electron acceptor 2,6-dichloroindophenol (DCIP) in enriched mitochondrial fractions. The complex II inhibitor TTFA was added at a concentration of 0.1 mM, and only the TTFA-sensitive activity is given here. Values represent the mean SQR activity in 2 independent small-scale mitochondrial preparations of mutant and wild-type like seedlings.

To measure complex II activity, mutant and wtl segregants were pooled for small-scale mitochondrial preparations. Of the different enzyme assays for complex II, succinate to ubiquinone reduction (SQR) using 2,6 dichloroindophenol as electron acceptor gave the most robust results in relatively impure mitochondrial preparations (León et al., 2007). The activity was measured before and after addition of the complex II inhibitor 2-thenoyltrifluoroacetone (TTFA) to distinguish complex II activity from other succinate reduction reactions. The TTFA-sensitive succinate turnover was similar in all three genotypes, i.e. *nfu4 nfu5* mutant, wtl segregants and true wild type (Fig. 7C). Thus, both aconitase and SDH had a normal operational capacity in *nfu4 nfu5* mutant seedlings, in contrast to the decrease in activities of these enzymes in yeast *nfu1*Δ mutants.

### NFU4 and NFU5 proteins interact with the mitochondrial lipoyl synthase LIP1

Previously, Arabidopsis NFU4 and NFU5 were shown to interact with the late-acting ISC proteins ISCA1a and ISCA1b (Azam et al., 2020a). To test if NFU4 and NFU5 can interact with downstream client Fe-S proteins, we carried out yeast 2-hybrid assays with mitochondrial lipoyl synthase (LIP1), biotin synthase (BIO2) and the major aconitase protein localized to mitochondria (ACO2). Biotin synthase is similar to lipoyl synthase in that an auxilary Fe-S cluster is destroyed to donate a sulfur atom in the last step of biotin synthesis, except that this cluster is a Fe_2_S_2_ cluster rather than a Fe_4_S_4_ cluster. Mitochondrial targeting sequences were removed from the coding sequences, which were cloned behind the activation domain (AD) or DNA binding domain (BD) of the Gal4 transcriptional activator. Yeast growth on medium without histidine indicated that the *HIS3* reporter gene is transcribed as a consequence of a direct protein interaction between NFU4/5 and LIP1 (Fig. 8A). The growth persisted under more stringent conditions with 3-amino-1,2,4-triazole (3AT), a competitive inhibitor of the *HIS3* gene product, suggesting that the interaction between NFU4, or NFU5, with LIP1 is relatively strong. However, no interactions between NFU4/5 and BIO2, nor with ACO2, were detected in these assays.

**Figure 8.**
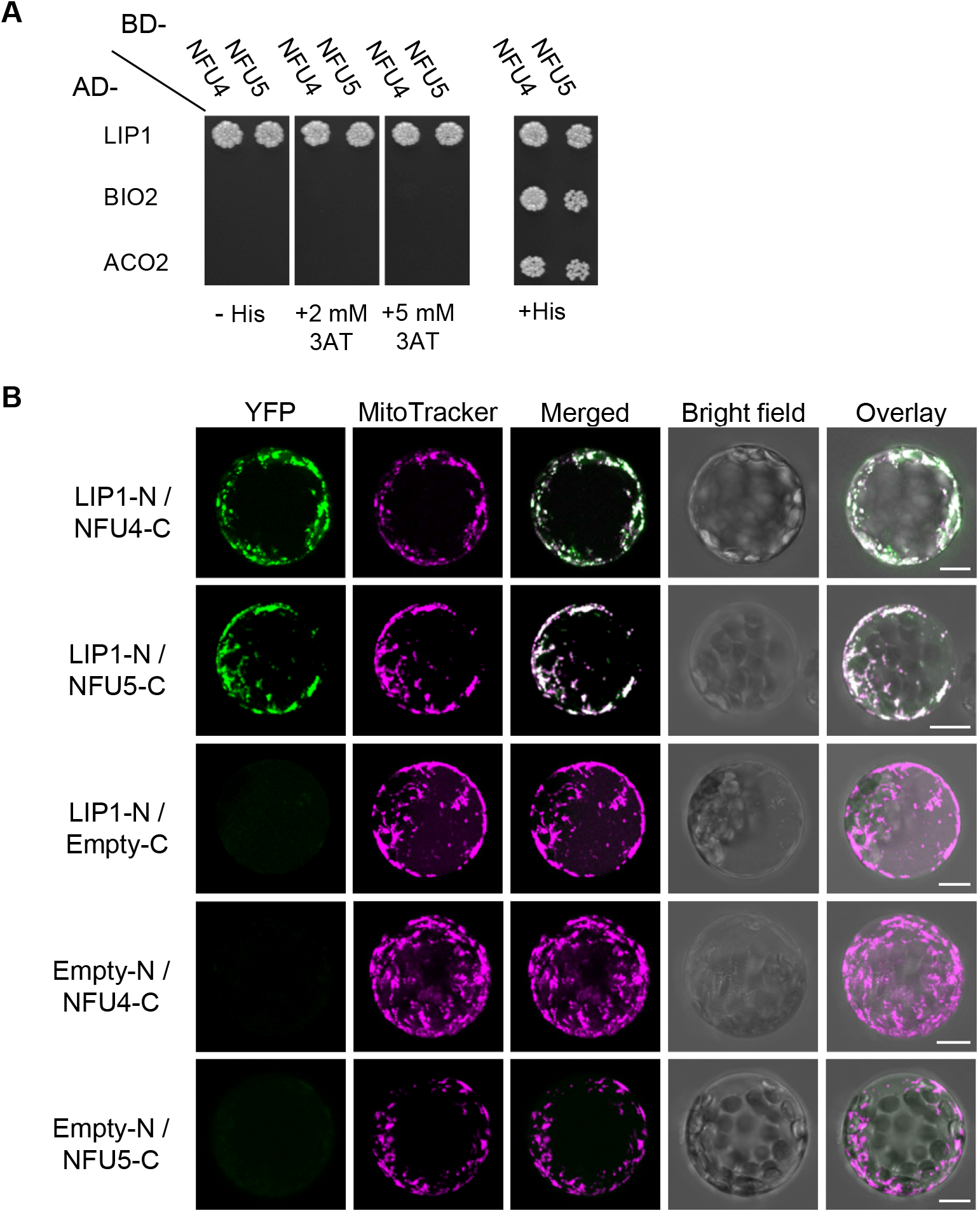
NFU4 and NFU5 proteins interact with LIP1. **A.** Yeast 2-hybrid analysis to test direct interaction between NFU4/NFU5 and mitochondrial lipoyl synthase (LIP1), biotin synthase (BIO2) and the main mitochondrial aconitase (ACO2). AD-, Gal4 activation domain; BD-, DNA binding domain, both at the N-terminal position. Images were taken after 5 days and are representative of at least 3 independent transformations. **B.** Bimolecular Fluorescence Complementation to test interaction between NFU4/NFU5 and LIP1. The coding sequences were placed upstream of the N-terminal or C-terminal region of YFP, and the plasmids transformed into Arabidopsis protoplasts. Results are representative of at least two independent transfection experiments and ≥20 fluorescent cells per transformation event. Images are provided with (Fig. 8B) and without (Supplemental Fig. S7) maximal Z-stack intensity projections. Scale bars: 10 μm.

To assess NFU4/5-LIP1 interactions in plant cells, we additionally performed bimolecular fluorescence complementation (BiFC) assays in Arabidopsis protoplasts using the LIP1 coding region fused upstream of the N-terminal domain of the YFP protein, and NFU proteins fused upstream of the C-terminal region of YFP (LIP1-N and NFU-C, respectively, in Fig. 8B and Supplemental Fig. S9). Positive BiFC signals indicated that LIP1 is in close proximity to NFU4 and NFU5 within plant cells, and that the LIP1 interaction with NFU4 or NFU5 occurs in the mitochondria as highlighted by the overlap of the YFP fluorescence with MitoTracker dye (Fig. 8B, merged panels). None of the proteins expressed alone could restore YFP fluorescence as previously shown (Azam et al., 2020a). Together, these results indicate that NFU4 and NFU5 interact equally well with LIP1 in the mitochondria, and likely assist with the insertion or repair of Fe-S clusters in LIP1.

## DISCUSSION

The physiological function of the mitochondrial NFU proteins has thus far not been studied in plants, except for one study in sweet potato which found that the gene encoding a mitochondrial NFU was upregulated under salt stress in a salt-tolerant variety (Wang et al., 2013). Here we show that the mitochondrial NFU proteins perform a key function in lipoate-dependent metabolism in the model plant Arabidopsis, most likely by providing Fe-S clusters to lipoyl synthase as shown for bacterial NfuA.

Arabidopsis has two paralogous genes, *NFU4* and *NFU5*, each encoding a functional mitochondrial NFU protein of relatively high abundance (Figs. 1 – 3). While no obvious growth phenotypes were observed in *nfu4* or *nfu5* single mutants, some of our findings suggest that Arabidopsis *NFU4* and *NFU5* may not be fully redundant. Single *nfu4* and *nfu5* mutants had impaired primary root growth in young seedlings (Supplemental Fig. S3). Moreover, during reproductive development, *nfu5* mutants could not be crossed with the *isu1-1* mutant as male or female parent, despite a functional copy of *NFU4* (Supplemental Fig. S4). Possibly, cell-specific expression of *NFU4* and *NFU5* in some specialized cell types, or differences in protein stability, could underlie these observations. Related to this is the question of whether NFU4 and NFU5 can form heterodimers or exist exclusively as homodimers. The yeast 2-hybrid assay testing for NFU4-NFU5 interaction was negative (Azam et al., 2020a), which is not due to technical issues since the same constructs showed positive interactions with other mitochondrial proteins such as ISCA1/2 (Azam et al., 2020a) and LIP1 (Fig. 8A). Thus, NFU4 and NFU5 are likely functioning as homodimers, with mostly overlapping but perhaps not identical physiological functions, which will be interesting to delineate in future studies.

The double knockout mutant of *NFU4* and *NFU5* in Arabidopsis is embryonic lethal (Fig. 3B–D), demonstrating that mitochondrial NFUs are essential in plants. The single *NFU1* homolog is also essential in mammals but the yeast gene is non-essential. How could this difference be explained? Lipoyl synthase is the main client Fe-S protein in all three types of organisms and an essential enzyme in higher eukaryotes, but not in yeast (Cronan, 2016). This is because lipoyl enzyme complexes can be bypassed, especially when yeast is grown on glucose and switches to fermentation. In bacteria, loss of lipoyl synthase (LipA) can be overcome by supplementation with lipoate in the medium (McCarthy and Booker, 2017), but this cannot rescue yeast *lip5* mutants (Sulo and Martin, 1993). Whether plant mutants in lipoyl synthase can be chemically complemented by external lipoate has not been investigated to our knowledge, but we found that addition of lipoate to the medium did not rescue the growth of *nfu4 nfu5* mutant seedlings in any way (Supplemental Fig. S10). The relatively high abundance of NFU4 and NFU5 compared to other Fe-S cluster maturation proteins would fit with a role of NFU proteins in reassembling the auxiliary cluster of LIP1, which is constantly turned over to insert two sulfurs into lipoyl cofactor. Particularly mitochondria in leaf mesophyll cells have a very high demand of lipoate for the H proteins of GDC.

To obtain *nfu4 nfu5* mutant lines with a clear phenotype, we used a strategy based on the transcriptional depletion of *NFU4* in a *nfu4 nfu5* genetic background (Fig. 5), after other approaches gave non-viable plants or no visible growth defects (e.g. Fig. 4). While the depletion approach was stringent and robust, the seedlings did not develop further than the 2-leaf stage, as reported for other essential genes (Rohde et al., 1999; Despres et al., 2001). This limited biochemical assays, especially since it was not possible to isolate mitochondria from the double mutant for enzyme assays. However, protein blot analysis showed a striking decrease in lipoylated proteins, especially the H proteins, and metabolite analysis provided further evidence of blocks in specific mitochondrial processes. While some variation between biological replicates of the mutant, for example in amino acid levels, could have been due to the precise timing of sampling relative to growth arrest, several highly significant changes were noted that have also been observed in other mutants in the lipoate biosynthesis pathway. Specifically, accumulation of glycine and elevated serine has so far been observed in all mutants affected in lipoyl cofactor biosynthesis and its octanoyl precursor (Ewald et al., 2014; Guan et al., 2017; Fu et al., 2020; Guan et al., 2020). Weaker mutant alleles (e.g. RNAi lines or non-essential genes) are primarily affected in the lipoylation of H proteins, whereas stronger alleles also showed decreased lipoylation of E2 subunits of PDH and KGDH. Interestingly, mutation of *GRXS15*, the mitochondrial glutaredoxin associated with the Fe-S cluster assembly pathway, also primarily affects lipoyl biosynthesis but not other Fe-S cluster-dependent processes (Ströher et al., 2016; Moseler et al., 2021). In-depth metabolomics analysis of a *grxs15* K83A mutant line showed accumulation of glycine, branched-chain amino acids and their keto-acids, a 5-fold increase in pyruvate but no significant difference in α-ketoglutarate (Moseler et al., 2021). Decreased lipoylation of the H proteins coincided with lower abundance of the polypeptides, as seen in *nfu4 nfu5* mutants (Fig. 6C).

*NFU1* mutations in human cells and yeast are known to cause defects in specific Fe-S enzymes, namely complex II and aconitase (Mayr et al., 2014; Melber et al., 2016). In Arabidopsis we saw no marked impairment of the activities of these enzymes in either single mutants or the *nfu4 nfu5* double mutant (Figs 4, S5 and 7). However, metabolite analysis of young leaves showed accumulation of succinate to similar levels in both *nfu4-2* and *nfu4-2 nfu5-*1 seedlings, suggesting that NFU4 may have a specific role in supporting succinate dehydrogenase function. Similarly, aconitase activity was slightly but significantly decreased in the double mutant compared to wild type when grown under standard conditions. By contrast, yeast 2-hybrid assays were negative for a protein interaction between NFU4 and ACO2 (Fig. 8). Interestingly, *in vitro*, ACO2 can receive an Fe-S cluster from NFU4 or NFU5 (Azam et al., 2020a), but this may simply reflect a possible thermodynamic transition, and not a physiological event. Alternatively, if ACO2 can receive its cluster directly from ISCA1/2 in vivo, this would bypass NFU4/5. It is also possible that the decreased aconitase and complex II activities represent an indirect effect of strong impairment of PDH. Further investigations are necessary to establish if NFU proteins do play a minor role in Fe-S cluster assembly on other proteins than lipoyl synthase.

The mitochondrial NFU proteins differ from their plastid counterparts in both structure and function. The plastid NFU proteins have an extra C-terminal domain which is a copy of NFU but lacking the CxxC motif. The function of this second domain is as yet unclear. Mutant analysis has shown that NFU1, NFU2 and NFU3 have partially overlapping functions required for the stability of Photosystem I and other plastid Fe-S proteins (Touraine et al., 2004; Yabe et al., 2004; Touraine et al., 2019; Berger et al., 2020; Roland et al., 2020). Interestingly, one of the proteins found to interact with NFU2 using yeast 2-hybrid and BiFC assays was the plastid isoform of lipoyl synthase LIP1p (Berger et al., 2020). The levels of lipoyl cofactor on the plastid PDH E2 subunits have not been analysed to date. Plastid PDH is an essential enzyme in fatty acid synthesis, and defects in this pathway have pleiotropic effects on photosynthesis (Bao et al., 2000; Ke et al., 2000).

In summary, our functional study of the plant mitochondrial NFU proteins reveals their importance for lipoyl cofactor biosynthesis and narrows down the position of these proteins in the downstream pathway of Fe-S cluster assembly.

## MATERIALS AND METHODS

### Plant material and growth conditions

The following T-DNA insertion mutants were obtained from Arabidopsis stock centres (ecotype in brackets, see Supplemental Table S1 for details): *isu1-1*, SALK_006332 (Col-0); *isu1-2*, GK_424D02 (Col-0); *nfu4-1*, SALK_035493 (Col-0); *nfu4-2*, SALK_061018 (Col-0); *nfu4-4*, SAIL_1233_C08 (Col-0); *nfu5-1*, WiscDsLoxHs069_06B (Col-0); *nfu5-2*, SK24394 (Col-4); *nfu5-3*, GT_3_2834 (Ler). Homozygous plants were selected using genotyping PCR and the T-DNA insertion site was verified by sequencing from the left T-DNA border. For *isu1-1* and *isu1-2*, the right border and contingent genomic DNA was also sequenced. Primers are listed in Supplemental Table S4. The *nfu5-1* allele needed to be outcrossed to remove an unrelated leaf phenotype.

Seeds were sown onto Levington’s F2 compost, or they were surface sterilised using chlorine gas and spread on 1/2-strength Murashige and Skoog (MS) medium containing 0.8% (w/v) agar. After vernalization for 2 days at 4°C, plants were grown under long-day conditions (16 hours light, 8 hours dark) at 22°C and light intensity of 180-200 μmol m^−2^ s^−1^ unless otherwise indicated. The generation and propagation of callus cell culture was performed as previously described (May and Leaver, 1993).

### Gene expression analysis

Total RNA was extracted using the Plant RNeasy kit (Qiagen), followed by DNase treatment (Turbo DNase kit, Agilent). The integrity of RNA in all samples was verified using agarose gels, and RNA purity was analysed by comparing 260/230 nm and 260/280 nm absorbance ratios (Nanodrop 2000, Thermo Fisher). RNA was quantified using a Qubit 2.0 fluorometer (Thermo Fisher). RNA (4 μg) was reverse transcribed to cDNA using Superscript III (Thermo Fisher). RT-qPCR reactions were made using SensiFAST master-mix (Bioline), in 20 μl volumes, each with 20 ng of cDNA. Reactions were measured in a Bio-Rad CFX-96 real-time PCR system and cycled as per the Bioline protocol. Data were analysed using the Bio-Rad CFX Manager 3.1 software, and were normalized using primer efficiency. The house-keeping gene *SAND* (*AT2G28390*) was used as reference gene (Han et al., 2013).

### Protein extraction, gel electrophoresis and protein blot analysis

To extract proteins, plant tissues were ground in cold buffer containing 50 mM Tris-HCl pH 8.0, 50 mM 2-mercaptoethanol and 1 mM phenylmethylsulfonyl fluoride, then centrifuged for 10 min to remove debris. Mitochondrial proteins from seedlings or cell culture were isolated according to (Sweetlove et al., 2007). Blue-native polyacrylamide gel electrophoresis (BN-PAGE) was carried out as described by (Wydro et al., 2013). For protein blot analysis, total protein extracts or purified mitochondria were separated by SDS-PAGE and transferred to nitrocellulose membrane by semi-dry electroblotting. To separate NFU4 and NFU5, 17% (w/v) acrylamide was used in the gels. Blots were labelled with antibodies diluted 1:5000 for anti-NFU4 and 1:2500 for anti-NFU5, followed by secondary anti-rabbit IgG conjugated to horseradish peroxidase, and detected by chemiluminescence.

To produce antibodies for NFU4 and NFU5, recombinant proteins were expressed and purified as described in (Zannini et al., 2018). These were used to raise polyclonal antibodies in rabbits by the Agro-Bio company (La Ferté Saint Aubin, France). Antibodies against the following proteins have been published: ISU1 (León et al., 2005); GRXS15 (Moseler et al., 2015); INDH and PDH E1 alpha (Wydro et al., 2013); and TOM40 (Carrie et al., 2009). Antibodies against aconitase (AS09 521) and GDC-H protein (AS05 074) were from Agrisera, Vännäs, Sweden; Antibodies against lipoate (ab58724) were purchased from Abcam, Cambridge, UK and monoclonal antibodies against actin (clone mAbGEa, product number MA1-744) were from ThermoFisher Scientific.

### Free amino acids and organic acids

Unbound, water soluble amino acids were extracted according to (Winter et al., 1992), with minor modifications. Approximately 30 mg of whole seedling were homogenised in 60 μl of 20 mM HEPES pH 7.0, 5 mM EDTA and 10 mM NaF. After addition of 250 μl of chloroform:methanol (1.5:3.5 volumes) and incubation on ice for 30 min, the water-soluble amino acids were extracted twice with 300 μl of HPLC-grade H_2_O. The aqueous phases were combined and stored at −80°C until further analysis. Samples were filtered and diluted 50x in water. Ten μl of each sample was derivatized using the AccQ tag kit following manufacturer’s instructions (Waters, UK) and 2 μl was used for injection. Derivatized amino acids were separated on a 100 mm × 2.1 mm, 2.7 μm Kinetex XB-C18 column (Phenomenex, USA) in an Acquity UPLC using a 20 min gradient of 1 to 20 % acetonitrile versus 0.1 % formic acid in water, run at 0.58 ml.min^−1^ at 25°C. The UPLC was attached to a TQS tandem mass spectrometer (Waters, UK) instrument for detection of the correct masses and quantitative measurement.

The organic acids citrate, α-ketoglutarate, malate, pyruvate and succinate were measured using a recently developed method (Marquis et al., 2017). In brief, 30 mg of whole seedlings were homogenised in 900 μl methanol:H_2_O (50:50 by volume, ice cold). After centrifugation at 15000 x *g* for 10 min at 4°C, the supernatant was evaporated in a GeneVac EZ-2 Plus (SP Industries,USA) and stored at −80°C until further analysis. Samples were then resuspended in 15 μl of water and derivatised with 50 μl of 10 mM 4-bromo-*N*-methylbenzymamine in acetonitrile and 25 μl of 1 M 1-ethyl-3-(3-dimethylaminopropyl) carbodiimide hydrochloride in acetonitrile:water (90:10 by volume) for 90 min at 37°C. Samples were run on an Acquity UPLC equipped with a Xevo TQS tandem mass spectrometer (Waters, UK) and a 100×2.1mm 2.6μm Kinetex EVO C18 column (Phenomenex). The following gradient of 1% (v/v) formic acid in acetonitrile versus 1% (v/v) formic acid in water, at 0.6 ml/min, 40°C was used: (0 min) 40%, (1 min) 40%, (2 min) 45%, (6.5 min) 95%, (7 min) 95%, (7.5 min) 100%, (8.5 min) 100%, (8.6 min) 40%, (12 min) 40%.

### Oxygen electrode and enzyme assays

Oxygen consumption rates of intact seedlings were measured using an Oxygraph (Hansatech, King’s Lynn, UK). Seedlings were transferred from ½ MS agar plates to 2 ml water in the oxygen electrode chamber and kept dark during the measurement. Aconitase activity was determined in total cell extracts in a coupled reaction with isocitrate, as described (Ströher et al., 2016). Succinate:ubiquinone reductase (SQR) activity of complex II, including preparation of mitochondria-enriched fractions from seedlings, was essentially as described by León et al (2007), except that 10 mM 2-thenoyltrifluoroacetone (TTFA) instead of 1 mM was used as specific complex II inhibitor. Addition of 10 mM TTFA in dimethyl sulfoxide (10 μl in a 1-ml reaction) inhibited SQR activity by 50-60%, but there was no inhibition with dimethyl sulfoxide alone.

### Yeast 2-hybrid assays

Open reading frames corresponding to the presumed mature forms of NFU4, NFU5, LIP1, BIO2, ACO2 (primers in Supplemental Table S4) were cloned into the pGADT7 and pGBKT7 vectors (Clontech) between the *Nde*I or *Nco*I and *BamH*I restriction sites in order to express N-terminal fusion proteins with the GAL4 DNA binding domain (BD) or with the GAL4 activation domain (AD), respectively, under the control of the *ADH1* promoter. The plasmids were co-transformed by heat shock treatment into the *GAL4*-based yeast 2-hybrid reporter strain CY306, which is deficient for cytosolic thioredoxins (MATα, *ura3-52*, *his3-200*, *ade2-101*, *lys2-801*, *leu2-3*, *leu2-112*, *trp11-901*, *gal4-542*, *gal80-538*, *lys2::UASGAl1-TATAGAL1-HIS3*, *URA3::UASGal4* 17MERS(x3)-TATACYC1-LacZ, *trx1::KanMX4*, *trx2::KanMX4* (Vignols et al., 2005). Transformants were selected on yeast nitrogen base (YNB) medium (0.7% yeast extract w/o amino acids, 2% glucose, 2% agar) without tryptophan and leucine (-Trp, -Leu). Two clones were selected and interactions were observed as cells growing on YNB medium in the absence of histidine (-His-Trp-Leu) at 30°C. The strength of the interactions was evaluated by challenging growth in the presence of 2 or 5 mM of the competitive inhibitor of *HIS3* gene product 3-amino-1,2,4-triazole. Images were taken 5 days after spotting on plates (7 μl/dot at an optical density of 0.05 at 600 nm). Absence of auto-activation capacities of the *HIS3* reporter gene by all constructs used in this study was systematically assayed after co-transformation of yeast cells by individual constructs producing a fusion protein with the corresponding empty pGADT7 or pGBKT7 vectors.

### Confocal microscopy

For localization of GFP-fusion proteins, full-length open reading frames (ORFs) of *NFU4* and *NFU5*, or the predicted MTS (amino acids 1-74 for NFU5 and 1-79 for NFU4) were amplified from *A. thaliana* leaf cDNA and cloned between *Nco*I and *BamH*I sites of pCK-GFP-C65T (2x *CaMV* 35S promoter, C-terminal GFP (Reichel et al., 1996). For BiFC, full-length ORFs of *NFU4*, *NFU5* and *LIP1* were amplified and cloned upstream of the C-terminal and N-terminal regions of the mVenus YFP protein into the pUC-SPY**C**E and pUC-SPY**N**E vectors (Walter et al., 2004, abbreviated -C and -N in figures, respectively) using the restriction sites *Xba*I and *Xho*I or *Sma*I. Primers for amplification are listed in Supplemental Table S4.

Leaf protoplasts were prepared from 21- to 28-day old *Arabidopsis* seedlings, transfected according to (Yoo et al., 2007) and imaged after 20 - 24 h. Prior to confocal analyses, the protoplasts were incubated with 20 nM MitoTracker® Orange CMXRos (ThermoFisher) to fluorescently label the mitochondria.

Image acquisition for NFU-GFP localization was performed using a Zeiss LSM780 confocal laser scanning microscope, and a water-corrected C-Apochromat 40× objective with a numerical aperture (NA) of 1.2. Signals were detected according to the following excitation/emission wavelengths: GFP (488 nm/494–534 nm), MitoTracker (561 nm/569–613 nm) and chloroplast autofluorescence (488 and 561 nm/671–720 nm). Pictures were analyzed using ImageJ software (https://imagej.nih.gov/ij/).

Image acquisition for BiFC was performed on a Leica TCS SP8 confocal laser-scanning microscope using a water x40 objective. Wavelengths for excitation/emission were: mVenus (514 nm/520–550 nm), MitoTracker (560 nm/580-620). Images were obtained using LAS X software and processed with Adobe Photoshop CS3 at high resolution. All confocal images shown here were captured without Z-stack intensity projection.

### Accession Numbers

Accession numbers are as follows: AT3G20970, NFU4; AT1G51390, NFU5; AT4G22220, ISU1; AT2G20860, LIP1.

## ACKNOWLEDGMENTS

We thank Brigitte Touraine, Cyril Magno and Frédéric Gaymard (INRA Montpellier) for mutant selection, seed propagation and initial phenotyping; Delphine Bernard (Department of Plant Sciences, University of Cambridge) for initial studies on the *isu1-1* and *nfu4-2* mutants; Rob Green and Lucy Anderson (John Innes Centre, JIC) for technical assistance; Markus Dräger (JIC) for root growth measurements and cloning of the ABI3:NFU4 construct; Baldeep Kular (JIC Metabolomics Platform) for amino acid analysis; and the Montpellier PHIV Platform for assistance with confocal microscopy.

